# Dissecting the role of non-coding RNAs in the accumulation of amyloid and tau neuropathologies in Alzheimer’s disease

**DOI:** 10.1101/095067

**Authors:** Ellis Patrick, Sathyapriya Rajagopal, Hon-Kit Andus Wong, Cristin McCabe, Jishu Xu, Anna Tang, Selina H. Imboywa, Julie A. Schneider, Nathalie Pochet, Anna M. Krichevsky, Lori B. Chibnik, David A. Bennett, Philip L. De Jager

## Abstract

**Background:** Given multiple studies of brain microRNA (miRNA) in relation to Alzheimer’s disease (AD) with few consistent results and the heterogeneity of this disease, the objective of this study was to explore their mechanism by evaluating their relation to different elements of Alzheimer’s disease pathology, confounding factors and mRNA expression data from the same subjects in the same brain region.

**Results:** We report analyses of expression profiling of miRNA (n=700 subjects) and lincRNA (n=540 subjects) from the dorsolateral prefrontal cortex of individuals participating in two longitudinal cohort studies of aging. Evaluating well-established (miR-132, miR-129), we confirm their association with pathologic AD in our dataset, and then characterize their in disease role in terms of neuritic β-amyloid plaques and neurofibrillary tangle pathology. Additionally, we identify one new miRNA (miR-99) and four lincRNA that are associated with these traits. Many other previously reported associations of microRNA with AD are associated with the confounders quantified in our longitudinal cohort. Finally, by performing analyses integrating both miRNA and RNA sequence data from the same individuals (525 samples), we characterize the impact of AD associated miRNA on human brain expression: we show that the effects of miR-132 and miR-129-5b converge on certain genes such as EP300 and find a role for miR200 and its target genes in AD using an integrated miRNA/mRNA analysis.

**Conclusions:** Overall, miRNAs play a modest role in human AD, but we observe robust evidence that a small number of miRNAs are responsible for specific alterations in the cortical transcriptome that are associated with AD.

## Introduction

Late onset Alzheimer’s disease (AD) is an age-dependent neurodegenerative disorder characterized clinically by cognitive decline and pathologically by the accumulation of neuritic β-amyloid plaques (NP) and neurofibrillary tangles (NFT) in the brain. Currently genetic [1], epigenomic [2] and transcriptomic studies [3, 4] coupled with advances in imaging techniques [5, 6] have begun to sketch the sequence of events in the causal chain linking risk factors to a syndromic diagnosis of AD dementia. One of these events may be the dysregulation of gene expression by alterations in the expression of microRNA (miRNA) and long intergenic non-coding RNA (lincRNA) molecules.

miRNA are a class of small regulatory RNA that modulate gene expression via a multiprotein complex which facilitates the interaction between an miRNA and its complementary elements in the 3’UTR of target mRNAs to initiate transcript degradation and repression of protein production [7, 8]. Aberrant expression of miRNA and/or its target mRNAs have been implicated in abnormal neuron function [9] and in several neurodegenerative disorders [10,11]. Recently, certain miRNA, such as miR-132 have been associated with pathologic AD [12–16]. However, these studies were conducted in a modest number of subjects with limited phenotypic information, and few results are consistent across these studies [17].

lincRNA are RNA that are longer than 200 nucleotides and do not code for proteins[18]. As with most long non-coding RNA and unlike miRNA there is no clear common functional mechanism for lincRNA [18]. There is still debate over what percentage of lincRNA may be functional at all [19]. However, focusing on the lincRNA that lie in the same locus as protein coding genes may provide insight into their functional correlates.

We first evaluated the role of known miRNAs with measures of both neuritic plaque (NP) and neurofibrillary tangles (NFT) since they are key neuropathologic features of AD and allow us to explore the mechanism by which an AD-associated non-coding RNA contributes to disease. We secondarily expanded this effort to evaluate other miRNAs and lincRNAs to discover new associations. In addition, we leveraged transcriptome-wide RNA sequencing profiles, generated from the same RNA samples that were used to generate the miRNA profiles, to identify the functional consequences of altered miRNA expression on protein-coding genes in the human dorsolateral prefrontal cortex (DLPFC) and to identify additional cases where miRNA and mRNA AD associations converge in the human cortex.

## Results

### Demographic features and nature of our ncRNA data

The NanoString nCounter assay was used to measure miRNA expression from the DLPFC of each subject. At the conclusion of a rigorous quality control and preprocessing pipeline, expression measures from 309 miRNAs in 700 subjects were retained for downstream analysis. The subjects profiled in this study are participants in one of two prospective cohort studies of aging, the Religious Order Study (ROS) and Memory and Aging Project (MAP) which are designed to be merged for joint analyses [20, 21]. These studies enroll non-demented individuals and include detailed, annual ante-mortem characterization of each subject’s cognitive status as well as prospective brain collection and a structured neuropathologic examination at the time of death. The study design of ROS and MAP yields an autopsy sample that includes a range of syndromic diagnoses and neuropathologic findings that are common in the older population. While this is not a true population-based sample, it captures the diversity of the older population at the time of death and is different than the collection of subjects typically used for age-matched case-control studies [12, 13,15, 16]. The subjects in the study have an average age of 88, 61% meet criteria for pathologic AD by NIA Reagan criteria [22] and 64% are female. To evaluate the nature of our data in relation to published results, we first assessed whether the expression of miRNAs are associated with a pathological diagnosis of AD according to the NIA Reagan criteria [22], focusing on an evaluation of previously reported miRNA associations [12, 13, 15,16].

### miRNA associated with a pathologic diagnosis of AD

We used a linear regression model controlling for age, sex, study, proportion of neuronal cells, postmortem interval and a measure of RNA quality (RNA integrity (RIN) score) to evaluate associations of miRNA with pathologic AD. In this analysis of the human DLPFC, two previously reported associated miRNAs are significantly associated with pathologic AD, exceeding a threshold defined using a Bonferroni correction for the number of miRNA tested, p< 1.6×10^-4^ (**Table 1, Figure 1**). Specifically, miR-132 (P = 2×10^-8^) and miR-129-5p (P = 3.9×10^-6^) were found to be diminished in expression in subjects with AD, consistent with prior studies [12, 13]. Thus, our data confirm well-validated miRNA associations with AD. To provide perspective on effect size, miR-132 explains 6.7% of the variance in AD, which is similar to the 6.1 % variance explained by *APOE* ε4 in our data and 6% in previous literature [23]. In contrast, the miRNA effect is more than the <1% of variance explained by genetic susceptibility variants in *ABCA7, CD33,* and *CR1* that were originally discovered in genome-wide association studies [23].

**Table 1.** miRNA and lincRNA associated with AD, NP or NFT.

|  | *coefficient: AD* | *p-value: AD* | *coefficient: NP* | *p-value: NP* | *coefficient: NFT* | *p-value: NFT* |
| --- | --- | --- | --- | --- | --- | --- |
| *miR-132* | -0.26 | **2.00E-08** | -0.29 | **1.40E-11** | -0.31 | **4.10E-08** |
| *miR-129-5p* | -0.11 | **3.90E-06** | -0.15 | **6.90E-11** | -0.17 | **2.20E-08** |
| *miR-129-3p* | -0.095 | 0.0013 | -0.13 | **1.80E-06** | -0.12 | 0.00085 |
| *miR-99b* | 0.081 | 0.0034 | 0.075 | 0.0037 | 0.14 | **6.00E-05** |
| *linc-CTSD-3* | -19 | 0.0012 | -22 | **6.90E-05** | -33 | **8.60E-06** |
| *linc-BRD9-1* | 32 | 0.00025 | 35 | **2.40E-05** | 23 | 0.036 |
| *linc-ADC* | 3.8 | 0.18 | 3.8 | 0.15 | 14 | **6.70E-05** |
| *linc-RNFT2-1* | 3.3 | 0.36 | 3.4 | 0.31 | 18 | **7.10E-05** |

**Figure 1.**
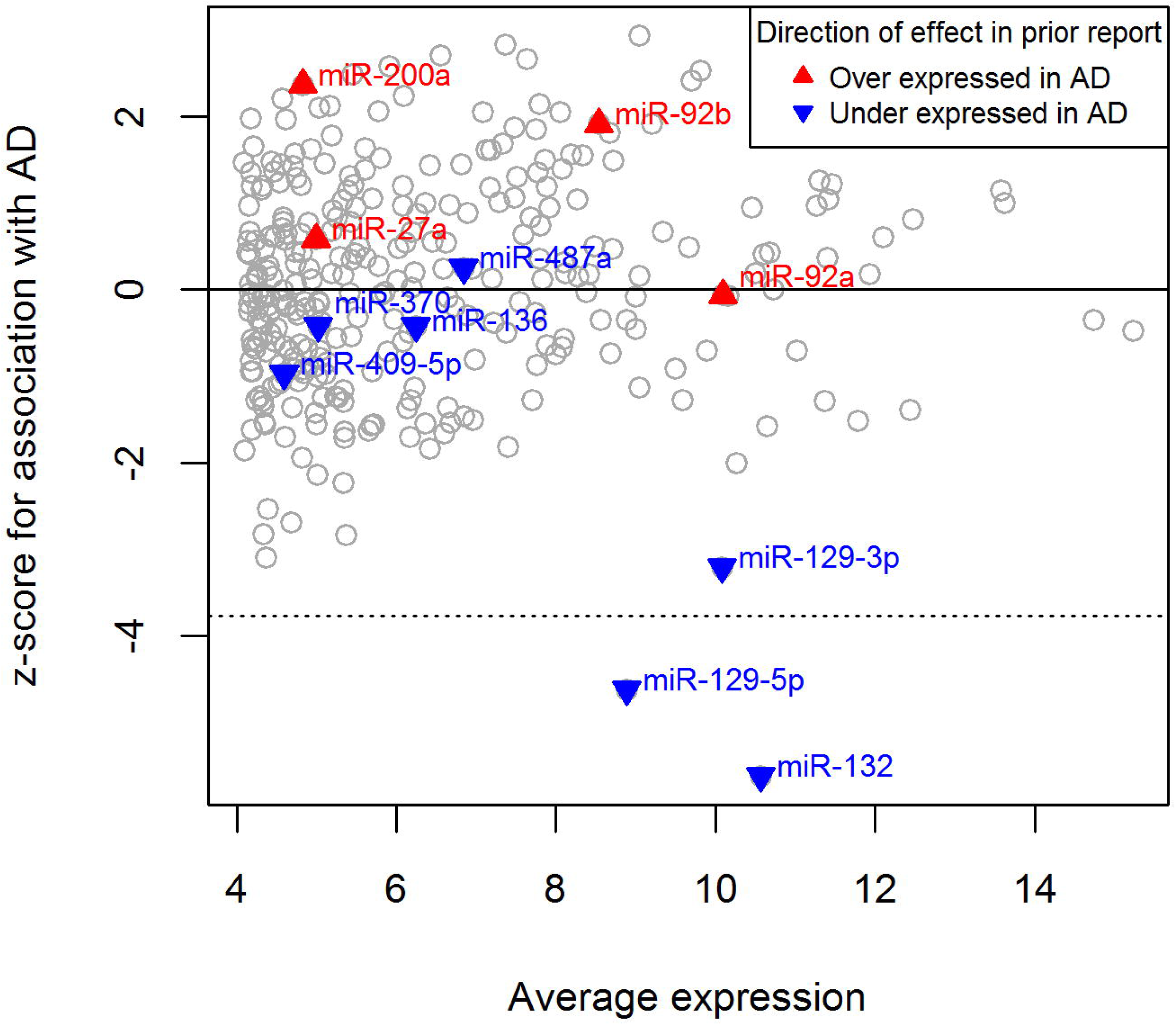
Validation of miRNAs associated with AD in other studies. The relationship between average miRNA expression (x-axis) and association with pathological AD in ROSMAP (y-axis) is plotted for each miRNA. The association between a miRNA and AD is calculated as a z-score after correcting for other factors; a negative z-score denotes decreased miRNA expression in the context of AD. Previously reported miRNAs [12] are highlighted by colored triangles, with red denoting higher expression in AD and blue reduced expression in AD in the original publication. The dotted horizontal line marks the z-score threshold of significance that is equivalent to a Bonferroni p-value cut-off of 0.05; none of the miRNAs with positively correlated expression are significantly associated with AD. The gray circles represent all other miRNAs tested in our study.

We next systematically reviewed our results for those miRNA that have previously been associated with AD in various brain regions [13,15,16]: they are catalogued in **Table 2. Figure 1** illustrates the strength of the association of miRNAs with pathologic AD in our cohort, highlighting 11 miRNAs that were deemed to be significantly associated with AD in a prior study of moderate sample size (n = 49 subjects) [12]. Of the 11 miRNA previously reported to be associated with AD in the prefrontal cortex, only miR-132 and miR-129-5p replicated. In addition, miR-129-3p (P=0.0013), miR-200a (P=0.018), miR-30e (P=0.0048) and miR-100 (P=0.011), let-7i (P=0.045) and miR-185 (P=0.013) display suggestive association and may warrant further investigation. The Bonferroni threshold (p<0.00016) that we employed may be overly conservative, but it provides a useful way to distinguish the most robustly associated miRNA. Given our large sample size (n=700), we have good power to confirm results with small effect sizes: we have 80% power to detect an effect small enough to only explain 2.2% of the variance in relation to AD (which is much smaller than the effect of mi132 described above). Thus, if other miRNAs that we tested have a role in AD, they are more in line with the magnitude of effect found with AD SNPs.

**Table 2.**
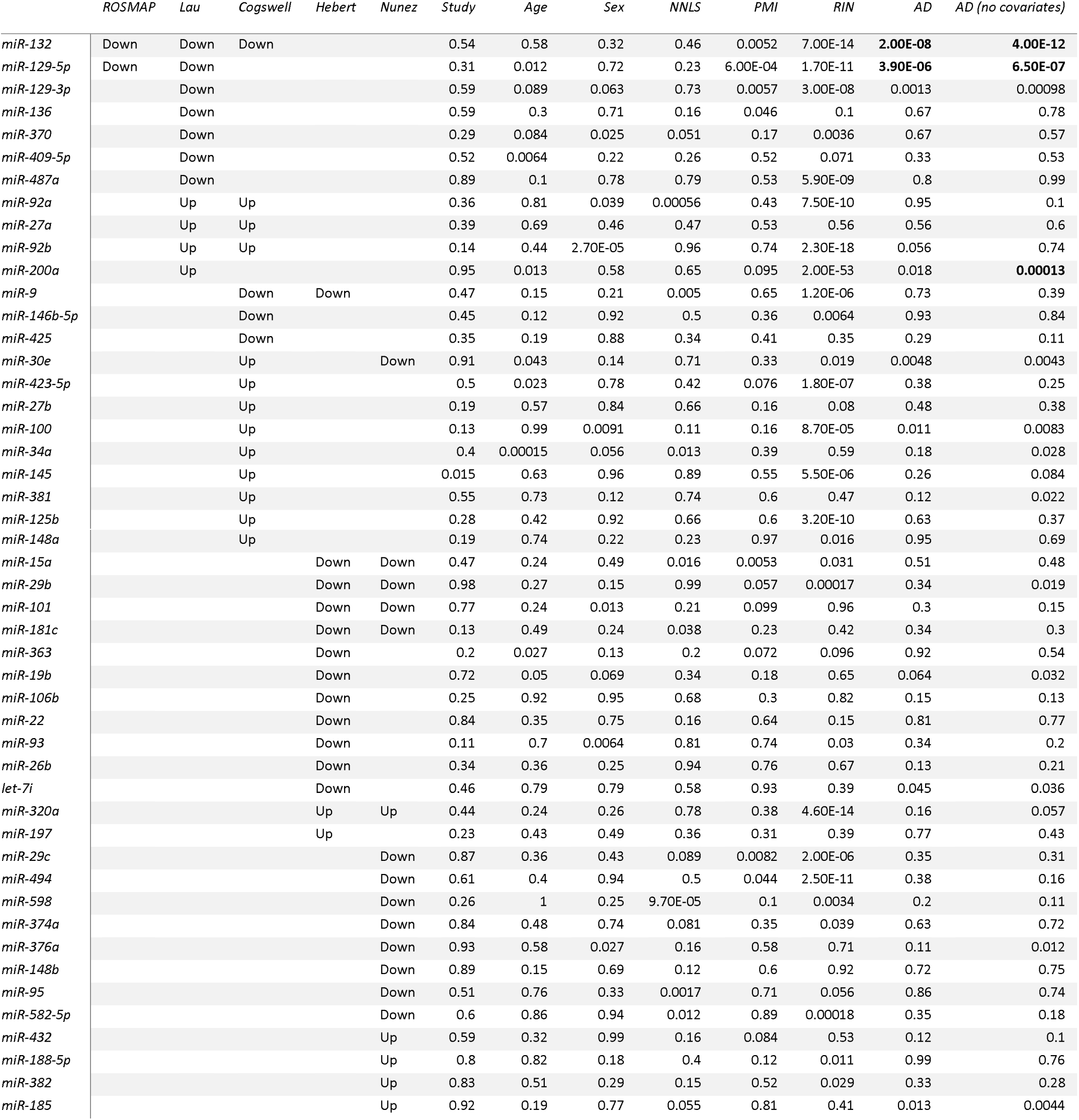
P-values from tests of association with AD and other covariates for miRNA implicated in other studies.

### The role of confounding variables in miRNA expression

There is a substantial lack of concordance in behavior between the previously reported AD miRNA signatures and their level of association in our cohort. While this could be explained, in part, by differences in the brain regions that have been assayed and the technologies used in the different studies, many of these miRNAs are highly associated with age, sex, the proportion of neurons in the profiled tissue sample, post-mortem interval and/or RIN score (**Table 2**). Thus, explicitly accounting for all of these factors is essential in brain miRNA studies, and the RIN score appears to play an especially significant role.

### Identifying miRNA associated with amyloid and Tau neuropathologies

Having described the nature of our data in relation to prior reports, we proceeded to our main goal of investigating the relationship of miRNA expression with specific features of AD neuropathology: namely, we investigated whether miRNA are associated with the accumulation of neuritic amyloid plaques (NP) and/or neurofibrillary tangles (NFT) that are defined by the presence of Tau accumulation. The pathologic AD-associated miRNA, miR-132 and miR-129-5p, were associated with both NP and NFT (**Table 1**). These two neuropathological features of AD are strongly correlated with one another, and it can be challenging to dissect the effect of one from the other unless large sample sizes are available. In addition, we found that, among the tested miRNAs, miR-129-3p (P = 1.8×10^-6^) was associated with NP, and miR-99b (P = 6×10^-5^) was associated with NFT, exceeding our significance threshold (p<0.00016).

### Dissecting the network of miRNA associations in AD

The complexity of the relationships between miRNA expression, the burden of NP and NFT neuropathologies, a pathologic diagnosis of AD, technical variables and demographic variables are demonstrated in **Figure 2a**. Here, we consider all variables in a single model to resolve the most likely primary association between each miRNA and the neuropathologic traits. These traits are all correlated with one another (**Figure 2d**), making it difficult to understand where an miRNA’s primary effect may be exerted simply by looking at the univariate results: for example, while miR-132 is significantly associated with both NP and NFT in separate analyses (**Table 1**), it appears, in the joint network model, that its effect may be driven predominantly through NP. This is visualized by the edge linking miR-132 and NP in the network model. The effect on NFT may be secondary and result from the accumulation of NP. To elaborate a more comprehensive network model, we included all of the miRNAs with either a significant or suggestive (p<0.05) association to one of the AD pathologies. Overall, in **Figure 2a** we see that the role of miRNAs in AD begins to be resolved, being linked with either NP or NFT. These associations with a pathologic features relating to either amyloid or tau pathology are important to generate hypotheses that can be tested in future studies, particularly in mouse models that may recapitulate only one of these pathologies. We also note that the confounding variables included in the model can have significant effects on disease-associated miRNAs, which are independent of the associations with AD and its neuropathologic features.

**Figure 2.**
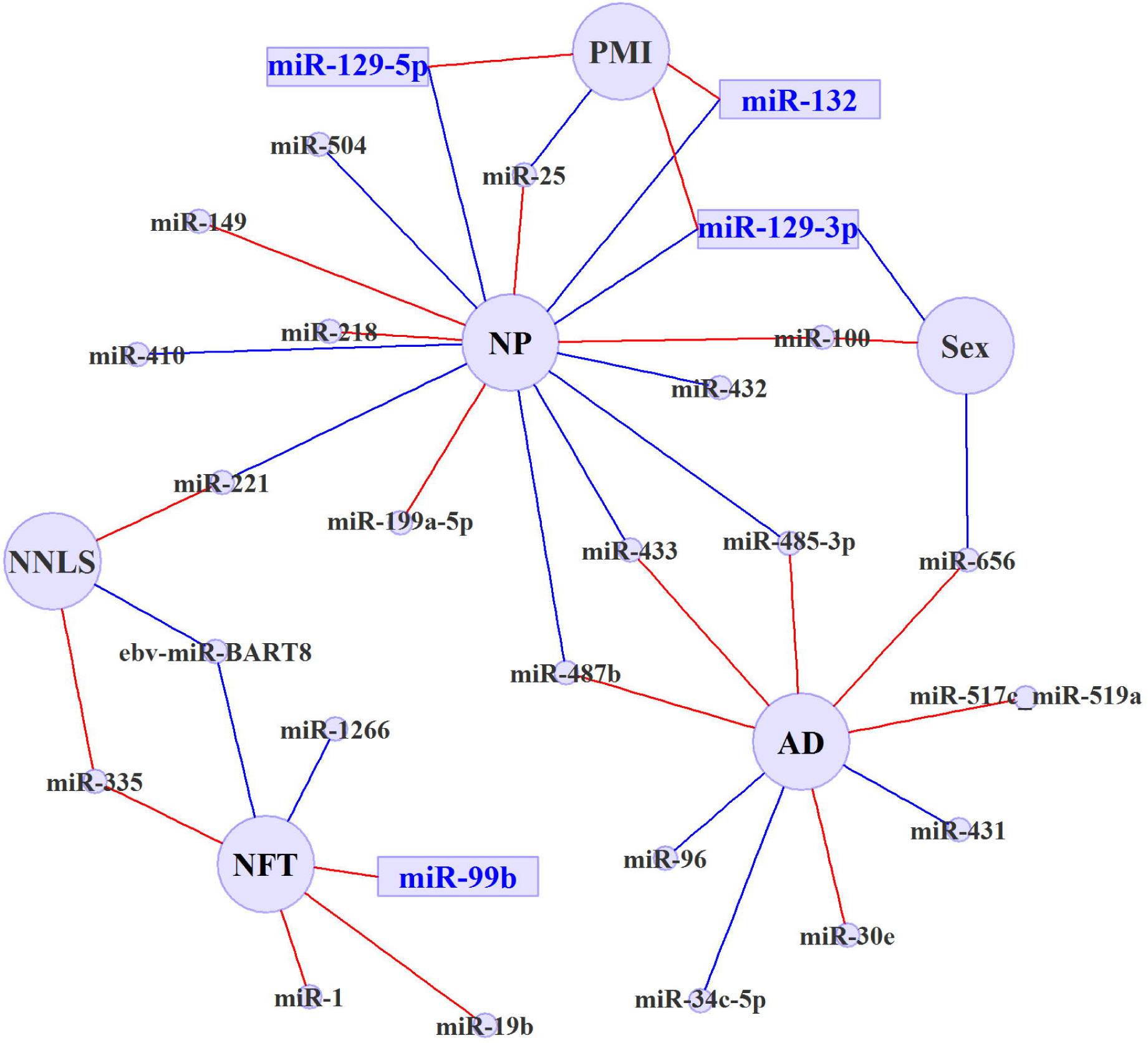
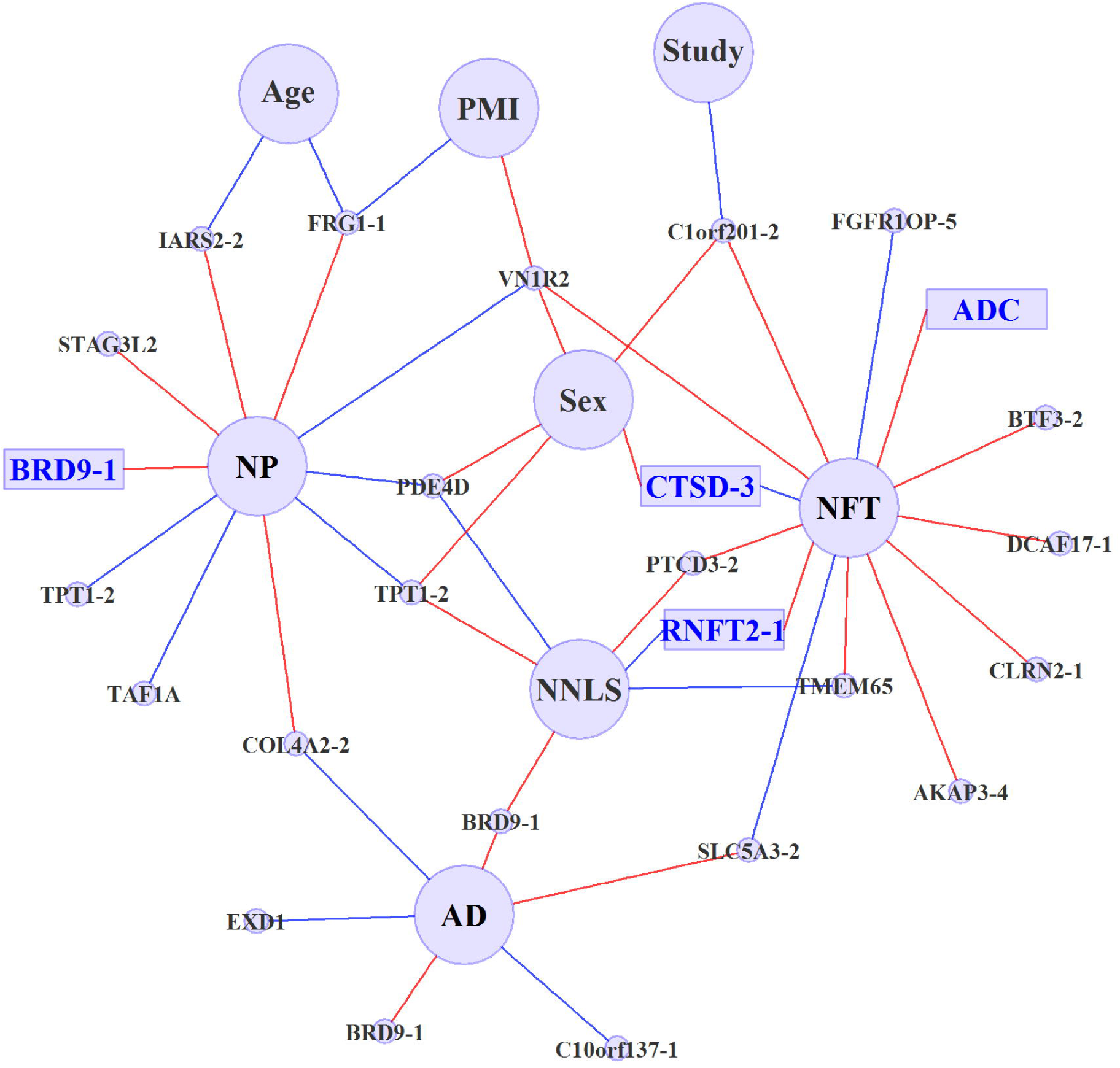
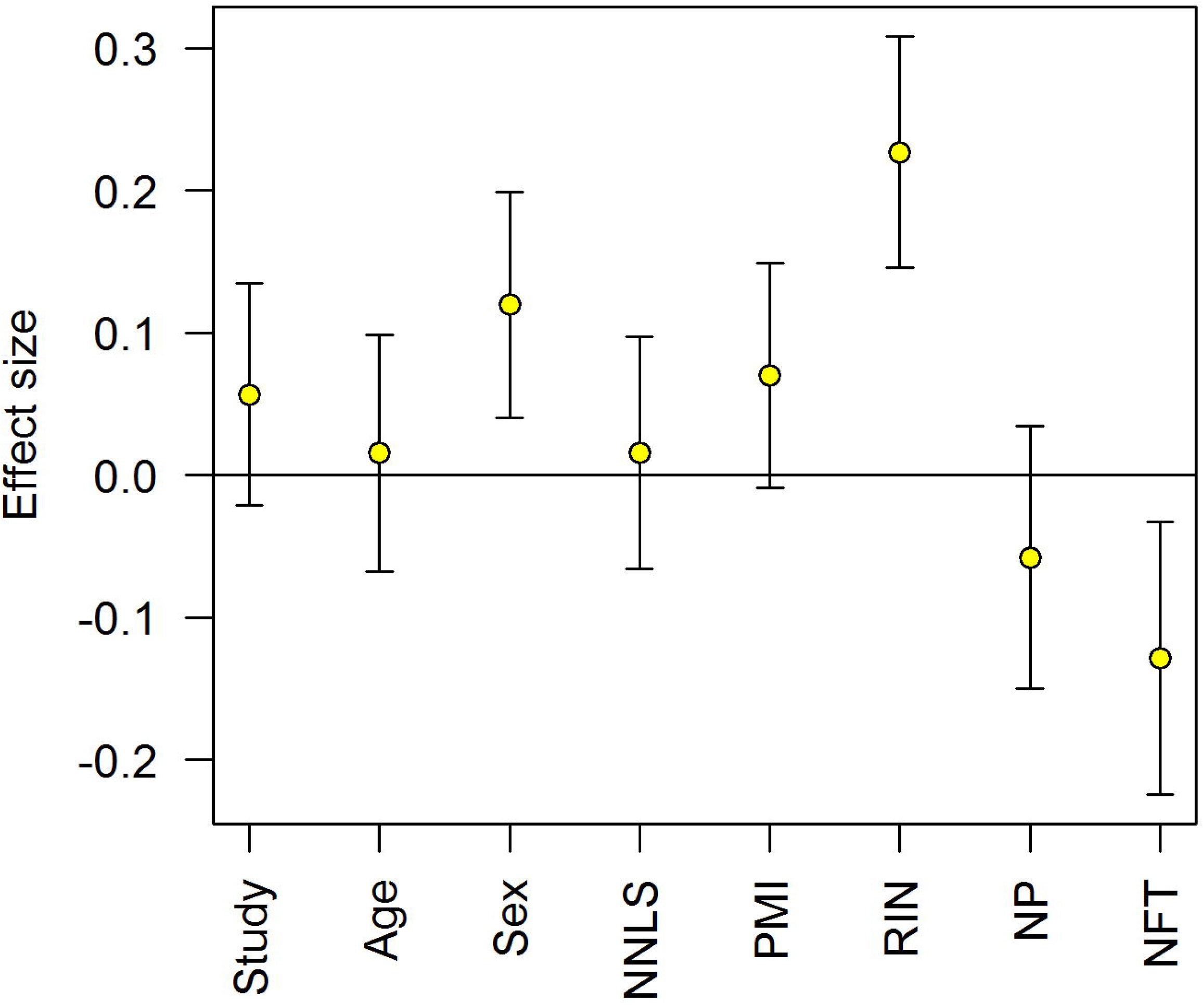
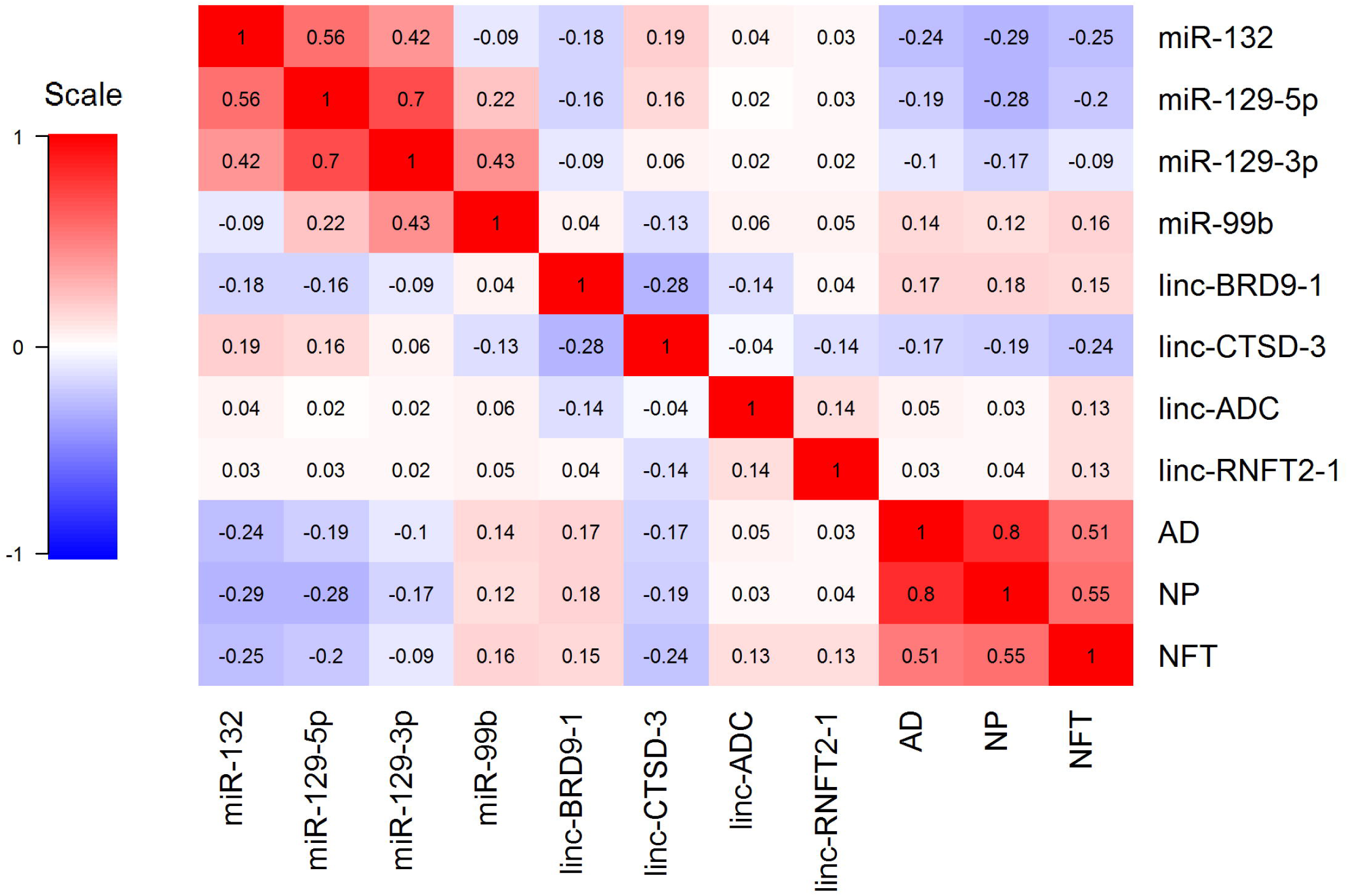
Network of miRNAs and lincRNAs associated with phenotypes. The relationships between (a) miRNA expression or (b) lincRNA expression and various demographic and technical features as well as neuropathologic outcomes are displayed in a network diagram. All miRNA and lincRNA with a significant or suggestive association are included in the network. These relationships were extracted via feature selection from linear models which explain an miRNA's expression level by using age at death (Age), neuronal composition (NNLS), sex, study of origin (Study, ROS or MAP), post-mortem interval (PMI), RNA integrity number (RIN), neuritic amyloid plaques (NP), neurofibrillary tangles (NFT) and pathological AD diagnosis (AD). Each node in the diagram represents either an miRNA or lincRNA (small circles) or a demographic or a neuropathologic variable (large circles). The significantly associated miRNA and lincRNA (**Table 1**) are represented by rectangles. A red edge represents a positive association between the non-coding RNA and trait; a blue edge denotes an inverse association. As most of the miRNAs are associated with RIN, the associations with RIN were removed from this figure for clarity. (c) The association of linc-CTSD-3 expression with each variable is plotted after fitting a model including all variables. The standardized effect size of each of these explanatory variables and 95% confidence intervals are shown. This illustrates the fact that non-coding RNAs significantly associated with AD pathology can be strongly influenced by confounding variables, but that these effects are independent. (d) The correlations between the expression levels of eight miRNA and lincRNA with significant associations in our study are shown, along with correlations with the outcome variables (NP, NFT and AD). Each of the correlations are displayed in each cell of the correlation map and is colored by the strength of correlation, red for positive associations and blue for negative associations.

### Evaluating the functional consequences of AD-associated miRNA in human brain

We next attempted to delineate the downstream effects of the expressed miRNAs by integrating the miRNA data with gene expression data from the same brain region and the same individuals. In fact, both the miRNA and RNA sequence data were generated from the same RNA sample. Our approach involved separate analyses of miRNA and mRNA data, with the mRNA analyses being focused on sets of genes that are co-expressed and are predicted to be a target of one of the tested miRNAs. Specifically, pMim [24] was used to identify sets of genes that (1) are predicted to be targeted by an miRNA associated with pathologic AD, (2) lie in a common biological pathway and (3) are also associated with pathologic AD themselves in terms of mRNA expression. **Figure 3a** outlines how pMim is used to construct and test “miR-pathways” where Targetscan v6.2 [25] was used for prediction of miRNA targets and the GO biological processes [26] were used for pathway annotation. All 309 miRNA and their corresponding miR-pathways (sets of genes that are targeted by the same miRNA and share a common biological process) were tested; a joint statistic summarizes and ranks the evidence for both the miRNA and mRNA analyses. The top eight miR-pathways in which both the miRNA and the mRNA are significantly associated with AD are presented in **Table 3**.

**Figure 3.**
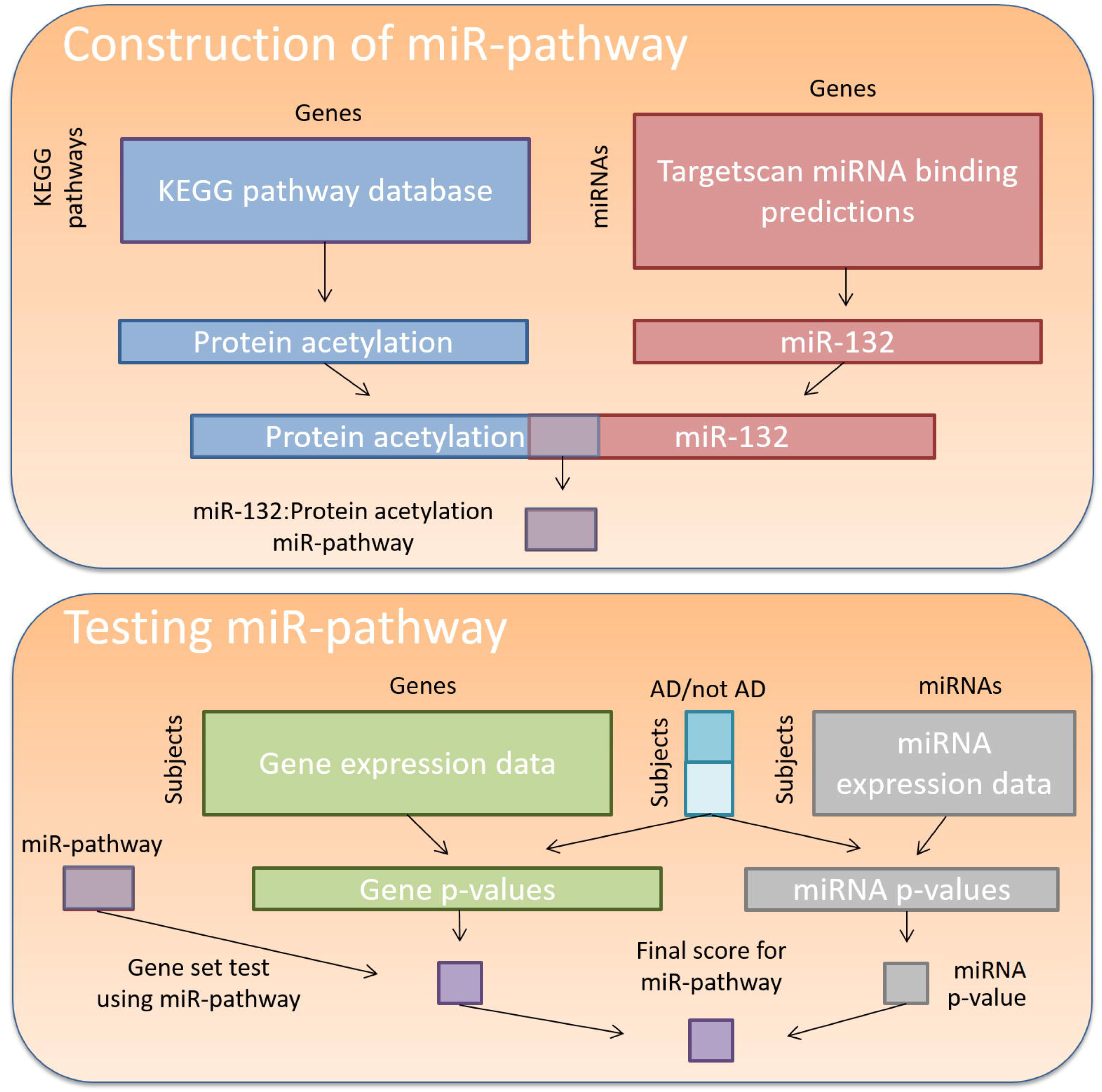
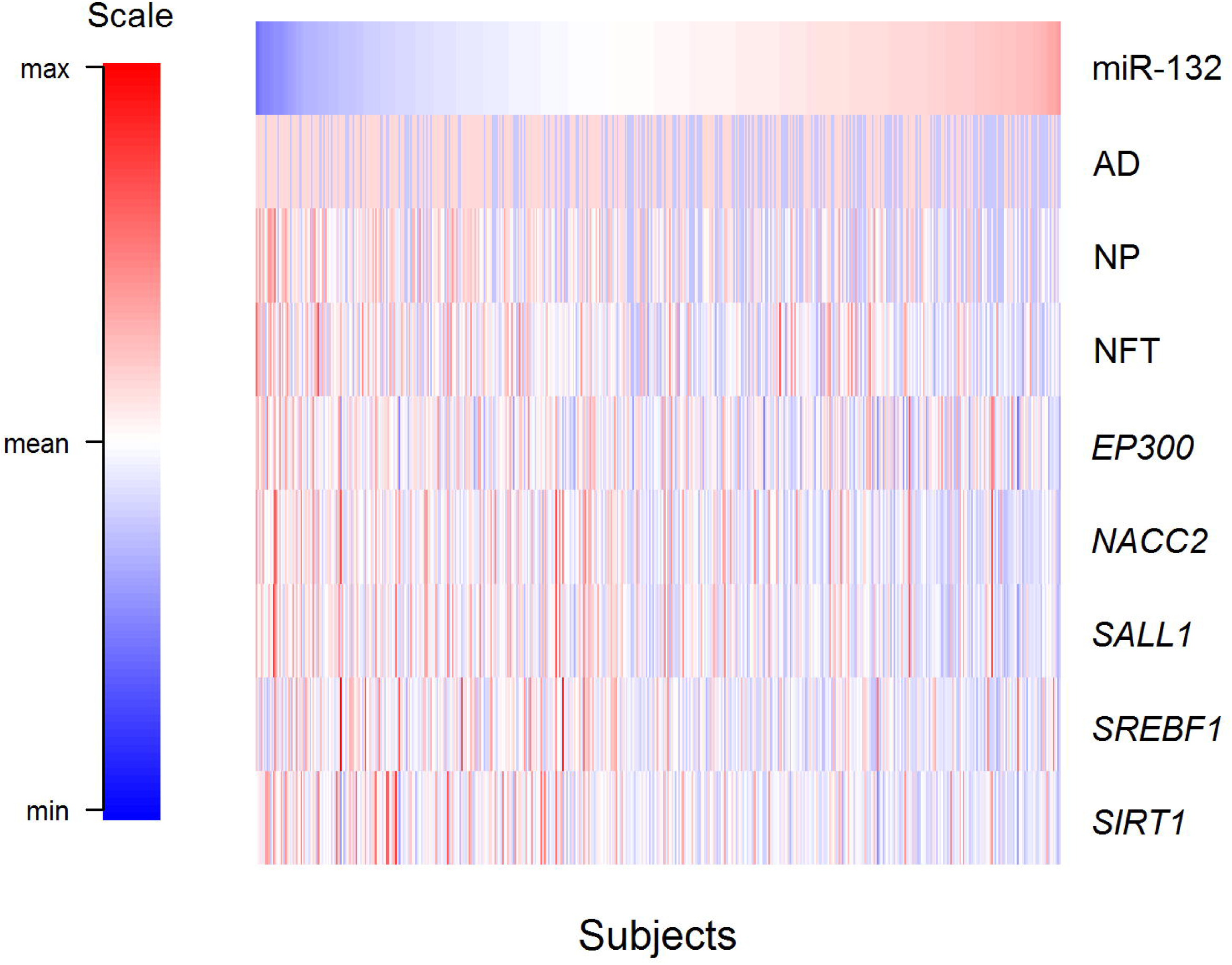
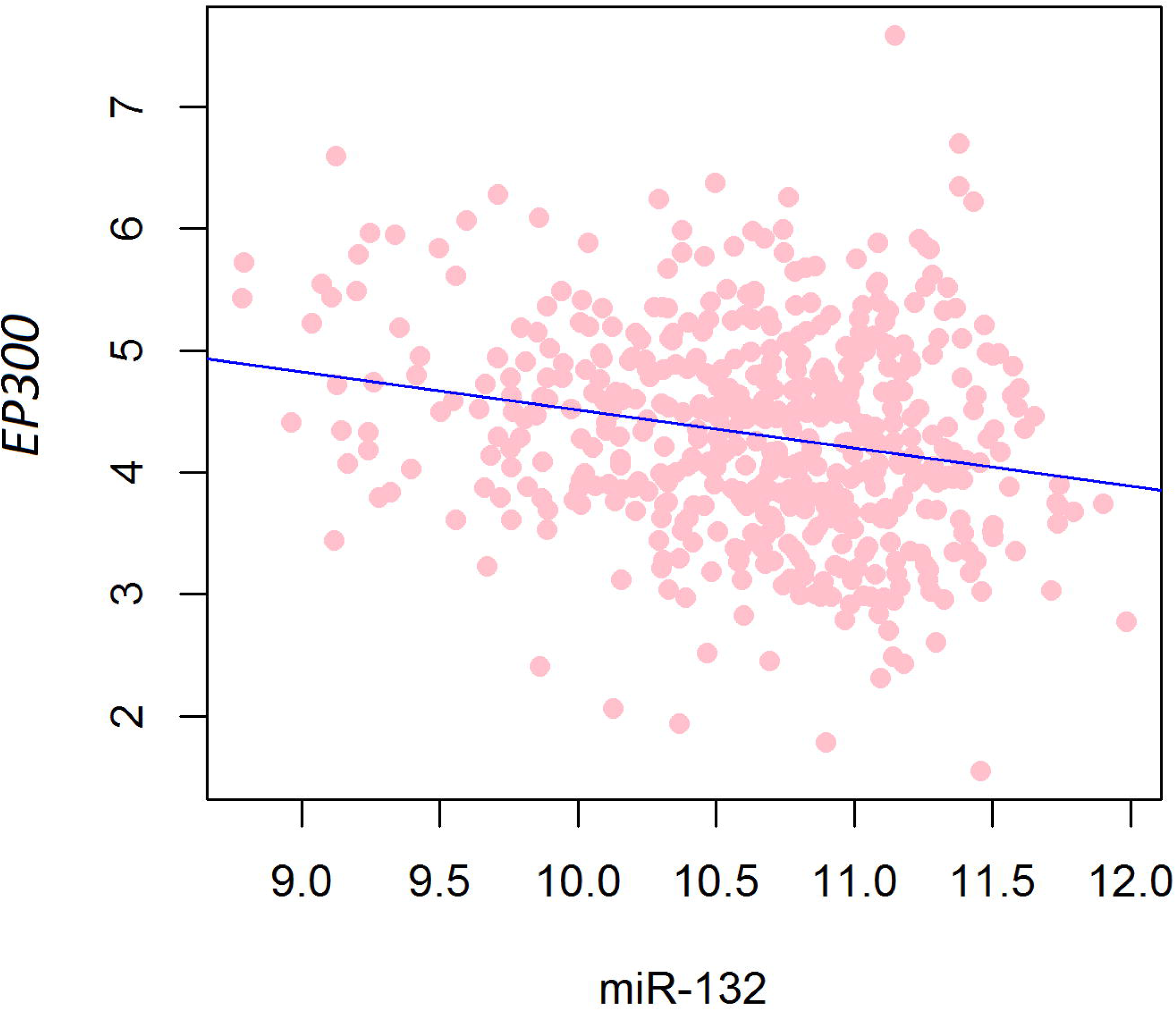
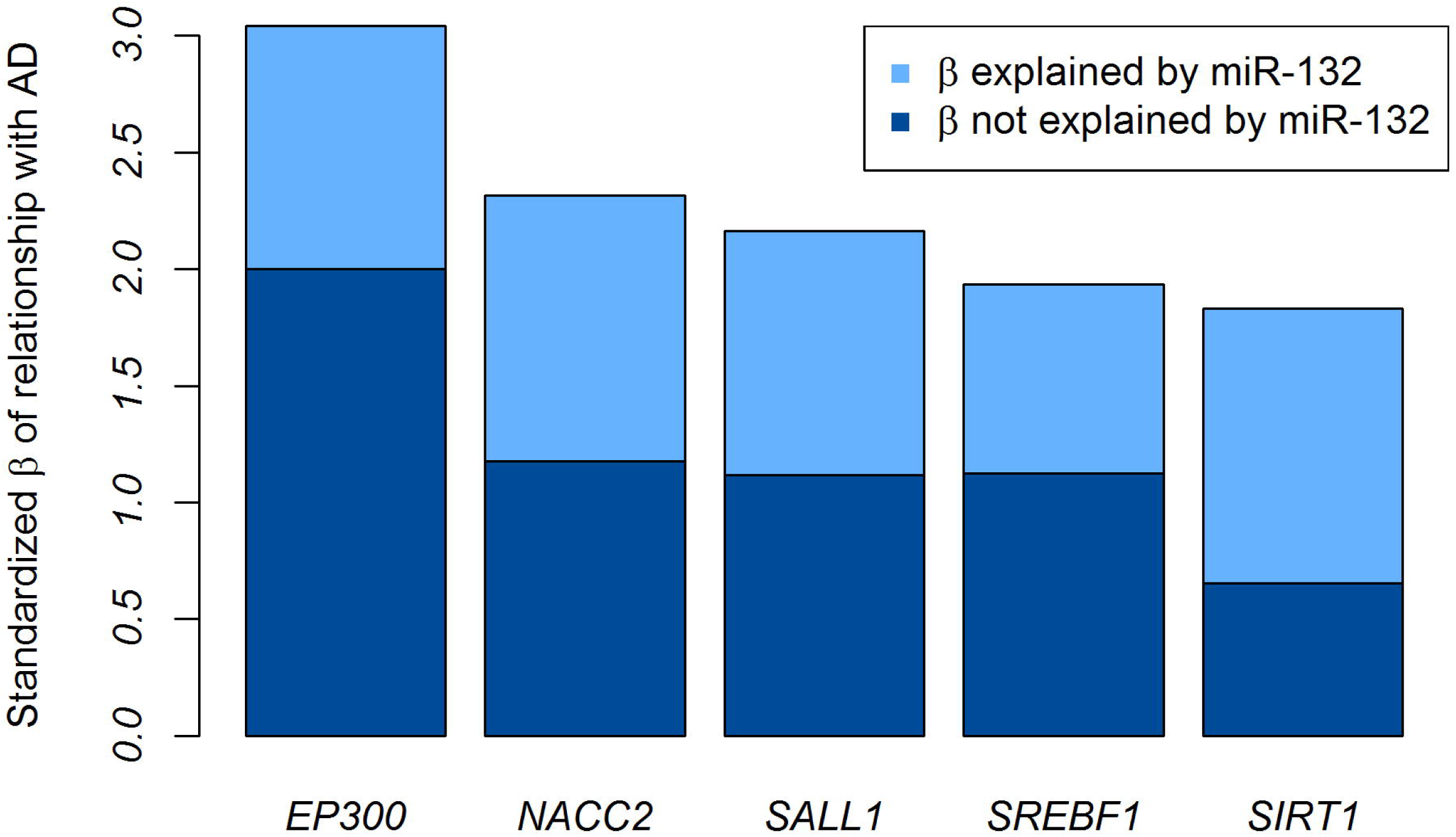
An integrated analysis of miRNA and their target mRNA. (a) This diagram presents an overview of (1) how the miR-132:Protein acetylation miR-pathway was constructed from the KEGG pathways database and Targetscan miRNA binding target prediction database and (2) how a score was derived to assess the association of this miR-pathway combination with pathological AD diagnosis. The approach is described in greater detail elsewhere [24]. (b) This heatmap illustrates the relation of miR-132 with the outcome measures (AD, NP, and NFT) and the mRNA expression levels of five of its target genes that are member of a protein acetylation miR-pathway. Each column presents data from one subject. Scaled expression values for a given variable are colored from red (to denote high expression) to blue (low expression). (c) The relationship between miR-132 and one of its targets, EP300, is demonstrated in a scatterplot. Each dot represents one subject. (d) The standardized effect sizes (β) capturing the strength of the relationship between the five protein acetylation genes and AD are plotted in this panel. We highlight the relative amount of the effect size (β) which can and cannot be explained by miR-132 expression, as miR-132 is predicted to be regulating each of these genes.

**Table 3.** Integrated analysis of miRNA and their target mRNA identifies miR:target combinations that are associated with AD.

| *miRNA* | *Pathways* | *Direction* | *Score* | *Genes in pathway targeted by miRNA* |
| --- | --- | --- | --- | --- |
| *miR-132* | protein deacetylation; protein deacylation; macromolecule deacylation | Down | 1.17E-09 | *EP300, NACC2, SALL1, SREBF1, SIRT1* |
| *miR-200a* | anion transport | Up | 1.07E-07 | *SLC20A1, SLC25A3, RIMS1, SLC35A1, SLC35D1, GLS, ATP8A2, ATP8A1, IRS2, SLC38A9, CXCL12, SLC16A7, SLC18A2* |
| *miR-129-5p* | regulation of transcription, DNA-templated | Down | 1.19E-07 | *EP300, NCOA1, ZBTB20, PBX3, LMNA, ZBTB5, ETS1, CHD7, CBX4, WWC2, YWHAB, CBX7, PKNOX1, ZBTB10, CBX6, PHIP, ZFY* |
| *miR-129-5p* | cell migration; cell motility; localization of cell | Down | 1.19E-07 | *SUN2, DLC1, LMNA, TMEM201, ETS1, RPS6KB1* |
| *miR-129-5p* | positive regulation of transcription, DNA-templated; positive regulation of transcription from RNA polymerase II promoter; positive regulation of RNA metabolic process; positive regulation of RNA biosynthetic process; positive regulation of nucleic acid-templated transcription | Down | 1.19E-07 | *EP300, NCOA1, LMNA, ETS1, PKNOX1, PHIP* |
| *miR-129-5p* | regulation of RNA metabolic process; regulation of nucleic acid-templated transcription; regulation of RNA biosynthetic process | Down | 1.19E-07 | *EP300, NCOA1, ZBTB20, PBX3, LMNA, ZBTB5, ETS1, CHD7, CBX4, WWC2, YWHAB, ACTN1, CBX7, PKNOX1, ZBTB10, CBX6, PHIP, ZFY* |
| *miR-129-5p* | cardiovascular system development; circulatory system development | Down | 1.19E-07 | *EP300, DLC1, LMNA, ETS1, CHD7, PKNOX1, TIPARP* |
| *miR-129-5p* | single-organism organelle organization | Down | 1.19E-07 | *SUN2, EP300, DLC1, NCOA1, LMNA, VAMP1, WHAB, SNX9, ACTN1, PAPD5, PHIP* |

Looking more closely at the top miR-pathways, we see that, in the DLPFC, miR-132 appears to be targeting a group of five genes (*EP300, NACC2, SALL1, SREBF1* and *SIRT1*) that are related to protein acetylation. All five of these protein acetylation genes lie within the top 20 predicted miR-132 target genes whose expression is inversely associated with miR-132 abundance in our cortical samples (p-values < 0.00018, **Table 4**). The miR-132 association with *EP300* expression confirms earlier reports and *in vitro* studies [12,14, 27, 28], illustrating the robustness of our results and the relevance of earlier studies. As shown in **Figure 3b-d**, miR-132 explains approximately a third of the association of each gene to AD, and therefore it does not appear to be the sole driver of the role of these protein acetylation genes in AD. While miR-132 may thus be influencing AD, in part, through histone acetylation and chromatin remodeling, miR-129-5p appears to be regulating genes related to the regulation of transcription. Interestingly, both miR-132 and miR-129-5p target *EP300*, which encodes the histone acetyltransferase protein E1A-associated cellular p300 transcriptional co-activator, and each miRNA explains some of the variation in *EP300* expression (adjusted R-squared 0.039 and 0.033 respectively). The role of these two miRNAs appears to be non-redundant: when they are considered together, they explain more of the variance in EP300 than either miRNA explains separately (adjusted R-squared 0.045), further illustrating the complex interactions involved in mRNA regulation by miRNAs.

**Table 4.** Predicted target genes that are negatively associated with miR-132 expression in the human cortex.

| *Gene* | *P-value: association with miR-132* | *P-value: association with AD* | *In protein acetylation pathway* |
| --- | --- | --- | --- |
| *CLMN* | 1.00E-14 | 0.0016 |  |
| *TJAP1* | 1.60E-12 | 0.00059 |  |
| *ANKRD40* | 9.30E-11 | 0.023 |  |
| *CXorf36* | 1.00E-10 | 0.0082 |  |
| *CDK19* | 2.60E-10 | 0.019 |  |
| *SSH2* | 6.10E-09 | 0.39 |  |
| *SEMA6A* | 4.90E-08 | 0.013 |  |
| *HIP1R* | 5.50E-08 | 0.0017 |  |
| *EP300* | 9.60E-08 | 0.0024 | ✓ |
| *DYNC1LI2* | 1.30E-07 | 0.023 |  |
| *NACC2* | 1.60E-07 | 0.021 | ✓ |
| *SOX5* | 1.60E-07 | 0.22 |  |
| *SIRT1* | 1.70E-07 | 0.069 | ✓ |
| *SALL1* | 5.10E-07 | 0.031 | ✓ |
| *MAPK3* | 4.10E-06 | 0.15 |  |
| *BCAN* | 2.00E-05 | 0.034 |  |
| *EPB41* | 0.00017 | 0.8 |  |
| *CTGF* | 0.00033 | 0.68 |  |
| *MEX3C* | 0.00038 | 0.53 |  |
| *SREBF1* | 0.00051 | 0.058 | ✓ |

In addition to these miR-pathways related to the validated miR-132 and miR-129-5p miRNAs, it is intriguing that miR-200a, which is suggestively associated to AD in our miRNA only analyses (**Table 2**), is the second strongest result in this pathway analysis because of the strong association of its downstream genes: this set of thirteen predicted target genes of miR-200a are all involved in anion transport. For these thirteen genes, an average of 18% of the mRNA association with AD is accounted by miR-200a. These integrated miRNA:mRNA analyses therefore report highly significant results drawn from autopsy tissue and set the stage for exploring specific downstream hypotheses; they also suggest that miRNAs are just one element contributing to the changes in gene expression found in AD since most of the variance of the affected genes – such EP300 and the miR200a targeted genes – remains unexplained at this time.

### Deregulation of lincRNA expression in relation to AD neuropathology

Our RNA sequencing data was also used to measure lincRNA expression from the DLPFC in the ROSMAP cohorts. At the conclusion of a rigorous quality control and preprocessing pipeline, expression measures from 454 lincRNAs in 540 subjects were retained for downstream analysis. The subjects with lincRNA expression are primarily a subset of the subjects with miRNA expression; 525 subjects have both miRNA and lincRNA expression data.

To investigate the relationships between lincRNA expression and the available neuropathologic measures, we used the strategy deployed in studying miRNA: we fit three independent linear models using age, sex, study, an estimate of the proportion of neuronal cells, post-mortem interval and RNA integrity number and either NP, NFT or pathological AD as covariates to explain lincRNA expression. The lincRNA that were significantly associated with NP, NFT or AD are shown in **Table 1**. No lincRNA is significantly associated with AD after Bonferroni correction. However, two are associated with NP (linc-CTSD-3 and linc-BRD9-1) and three with NFT (linc-CTSD-3, linc-ADC and linc-RNFT2-1); none of these lincRNAs have sequences that overlap the exons of protein coding genes.

We then repeated our network-building strategy to illustrate the primary associations among lincRNAs and the various demographic and pathological variables (**Figure 2b**). We use linc-CTSD-3 to illustrate the fact that the pathologic AD-associated non-coding RNAs can also be independently influenced by confounding variables, highlighting the need to include them in the modeling (**Figure 2b-c**). Like the miRNA network, no lincRNA maintains a relationship between both NP and NFT after feature selection.

The functional consequences of lincRNAs remain poorly understood, and we therefore could not repeat our pathway analysis. Since lincRNAs may influence the coding RNAs found in the same locus, we evaluated the association of the neighboring protein-coding genes with AD and the neuropathologic outcomes, but none of the coding transcripts were significantly associated (data not shown). While the previously reported BACE1-AS [29] was not in the lincRNA database used in this study, the positive association of BACE1 mRNA expression with AD was not replicated in our cohort (P = 0.63).

We have therefore investigated two different types of non-coding RNAs, miRNA and lincRNA, that may play a role in regulating gene expression in AD. Interestingly, while the expression level of different pathologic AD-associated miRNAs are fairly strongly correlated to one another, they are not strongly associated with lincRNAs, and the lincRNAs are only modestly correlated with one another (**Figure 2d**). This suggests that, while related, the molecular mechanisms to which these non-coding RNAs contribute in the context of AD may be largely different, with greater coordination among miRNA effects.

## Discussion

The number of individuals profiled in our cohorts and the prospective nature of the brain collection in these longitudinal studies of aging make our dataset a valuable resource for exploring, in greater detail, the role of miRNAs that have previously been associated with pathologic AD: our analysis meets a need for studies in larger datasets [17, 30] and offers a potentially less biased perspective of the disease than the comparison of AD cases and controls that are pulled from a brain bank to fit certain diagnostic criteria and were not collected prospectively. Aside from the two validated miRNAs (**Table 1**), we found suggestive evidence of association (p-values <0.05) for 8 of the 48 previously proposed pathologic AD-associated miRNAs, demonstrating that additional miRNAs may have smaller effects on AD. While all of the other studies age-matched their subjects, none explicitly modeled sex or other technical covariates in their analysis. In particular, we observe strong associations with RIN score, a technical measure capturing the quality of the RNA sample that led to spurious associations when not accounted for in the analysis. One study [12] reports the overall RIN scores for their hippocampal and prefrontal cortex samples; however, they do not provide a description of how this measure correlates with AD, and the incomplete RIN score data reported by another study [13] shows some potential differences between AD and controls. By accounting for the covariates that we have measured, we not only enhance confidence in the miRNA reported as being associated with AD pathology but also provide an opportunity to speculate on the reason why some miRNA may not have been validated due to the technical, clinical and demographic differences among subjects selected for different studies.

Our study has certain limitations, including the advanced average age at death (88 years) in these cohorts and the fact that they are representative of the older population but are not truly population-based, which limit the generalizability of our results. However, the high rate of autopsy (>90%) among study subjects ensures that our results are representative for the entire study population, which consists of subjects who are non-demented at the beginning of the study. Further, the data that we analyzed was generated from the cortex (gray matter) of the DLPFC. This is a practical compromise to attain large sample sizes, but it presents a challenge for future work as it is not clear which cell type may be driving a particular association. While accounting for the proportion of neurons in the tissue addresses some of the concerns that relate to the role of changes in the relative frequency of cell populations, future work in purified cell populations will be needed to resolve these questions more fully. The Nanostring technology used to measure miRNA expression could also introduce a technical bias in the replication of previously reported miRNA, and the use of probes to measure miRNA with Nanostring and of a predefined reference for aligning lincRNA data will limit the discovery of unannotated non-coding RNAs. Finally, we cannot comment on causality since we have a performed a cross-sectional analysis of brain tissue.

The availability of transcriptome-wide mRNA data from the same RNA samples in a large (n=540) subset of the ROSMAP subjects profiled for miRNA also provides a rare opportunity to directly explore the relation of miRNA with their putative target mRNAs and of miRNAs with a different class of noncoding RNAs, lincRNAs, in human tissue. Some of the lincRNAs are associated with AD pathology but their expression appears to be largely independent of the miRNAs. On the other hand, our large autopsy-derived mRNA sequence data has identified several different molecular pathways whose component genes have mRNA levels that are associated with AD and are targeted by AD-associated miRNA. The robustness of these results is nicely demonstrated by our lead miRNA, miR-132, which has been validated to be associated with AD in prior targeted studies [12,13, 31] and for which selected putative target genes have been evaluated in brain samples, including *EP300* and *SIRT1 [12,14]*, Here, we not only refine earlier observations by showing that the effect of miR-132 is mediated by the accumulation of amyloid pathology but also expand prior targeted studies of downstream effects by organizing the target genes in pathways to highlight cellular functions, such as protein acetylation that appear to be targeted by alterations in miR-132.

With our analyses, we have therefore begun to use autopsy data from a large set of well-characterized human subjects that capture the heterogeneity of older human brains to resolve which aspect of AD-related pathology is influenced by each miRNA of interest. The two well-validated miRNAs in AD illustrate this well: while miR-132 and mir-129-5p are strongly associated with the correlated amyloid and tau pathology measures, both miRNAs are more strongly associated with amyloid pathology than the accumulation of Tau pathology when the pathologies are included in the same model. This leads to very different experimental paths to further dissect the mechanism of these miRNAs in AD. In addition, using these quantitative pathologic traits that are more precise than a categorical diagnosis of AD, we find that some new miRNAs, such as miR-99b, that may have a stronger effect on a specific pathology, such as Tau/NFT. lincRNAs also appear to be involved, but the downstream consequences of these non-coding RNAs remain unclear. Nonetheless, the lincRNA associations bring another dimension to the broad narrative that emerges from our report: that molecular changes associated with AD include an important alteration in the regulation of cortical transcription, which is consistent with prior reports of epigenomic changes in certain model systems such as Drosophila Melanogaster DNA methylation profiles and the involvement of REST in AD [32, 33]. This narrative is also illustrated by miR-200 where the simple analysis of the miRNA alone is suggestive but not convincing of association with AD; however, an integrated analysis that also considers alterations of miR-200 target genes prioritizes this miRNA and transcriptional changes in anion transporters for further evaluation. Such integrated analyses of complementary data may be helpful to resolve the broader perspective on alterations in cellular function in AD. With this manuscript, we therefore provide a robust foundation of detailed neuropathologic associations that set the stage for a new generation of integrative analyses that considers different molecular measures generated from the same subjects and allows for the direct modeling of the complex phenotypic and molecular heterogeneity of the aging population at risk of AD and other dementia.

## Conclusions

By studying cortical levels of non-coding RNA in a large prospectively recruited cohort of well-characterized human subjects we had a unique opportunity to explore their associations with measures of both neuritic plaques and neurofibrillary tangles as well as the target mRNAs that they may be regulating. This work provides a robust foundation for future hypothesis-driven work to further dissect the mechanism of the reported associations to specific neuropathologic features of AD.

## Methods

Total RNA, including miRNA, was extracted from approximately 100mg sections of frozen postmortem brain tissue from the dorsolateral prefrontal cortex (BA 9/46) of subjects from two previously described longitudinal cohorts of aging, Religious Order Study (ROS) and Rush Memory Aging Project (MAP) [2, 34–36]. Tissue was thawed partially on ice and between 50-100mg of gray matter was dissected from the section then transferred immediately to 1mL of Trizol. The tissue was then quickly homogenized using the Qiagen TissueLyser and a 5mm stainless steel bead, for 30 seconds at 30Hz. The foam was settled with a quick spin, and the sample incubated for a minimum of 5 mins at room temperature. Debris was pelleted at 4°C at 12,000g for 10 min and Trizol was transferred to a new 1.5mL tube and proceeded as per Qiagen's miRNeasy Mini kit instructions, with volumes adjusted for 1mL instead of 700uL of Trizol until wash steps. RNA was eluted from the miRNeasy spin columns in 75uL of elution buffer, and quality tested by Nanodrop and Bioanalyzer RNA 6000 Nano Agilent chips. RNA yields averaged about 25ug and RIN scores ranged from 2-9 with an average of 6.5. RNA was normalized to 33ng/ul and plated into 96w plates for Nanostring processing using the nCounter Human miRNA Expression assay kit version 1 with reporter library file: NS_H_miR_1.2.rlf. The data collection from 733 post mortem brain samples was done at the Broad Institutes Genomics Platform (Broad Institute of Harvard and MIT, Cambridge, USA). Subjects from different diagnostic categories were distributed across experimental batches to reduce batch effects. To minimize variability at the ligation step, processing of the annealing and ligation steps was performed on the same thermocycler. Two thermocylcers were used for the purification steps, but all samples were placed in the same thermocycler for denaturation and hybridization steps. Two nCounter Prep Stations were utilized, but all samples were then scanned on the same Digital Analyzer. The data was collected in 8 batches of 96 samples and a single sample technical replicate was introduced as control in every single 96 well plates and sometimes twice in one single plate in two different cartridges.

### Quality control and dataset pre-processing

All data is available at https://www.svnapse.org/#!Svnapse:svn3219045. The miRNA from the Nanostring RCC files were re-annotated to match the definitions from the miRBase v19. The raw data from the Nanostring RCC files were accumulated and the probe-specific backgrounds were adjusted according to the Nanostring guidelines with the corrections provided with the probe sets. After correcting for the probe-specific backgrounds, a three-step filtering of miRNA and sample expressions was performed. First, miRNA that had more than 95% of missing expressions were removed. This is followed by removing samples that had more than 95% of miRNAs with missing expressions. Thus, the call-rates for the samples and the miRNA are set at 95%. Finally, all miRNA whose absolute value is less than 15 in at least 50% of the samples were removed to eliminate miRNA that had negligible expression in brain samples. After the miRNA and sample filtering, the dataset consisted of 309 miRNAs and 700 samples. A combination of quantile normalization and Combat [37], specifying the cartridges as batches for the miRNA data, was used to normalize the data sets. The strong association observed between miRNA expression and RNA-integrity was validated via qRT-PCR (data not shown) and verified not to be specific to the Nanostring platform.

The mRNA sequencing data have been described elsewhere [38]. Briefly, for the lincRNA analysis, a non-gapped aligner Bowtie [39] was used to align reads to a hg19 lincRNA reference [40] and then RSEM [41] applied to estimate expression levels for all transcripts in lincRNA resulting in a dataset with 454 measured lincRNA from 540 samples. A combination of quantile normalization and Combat [37] was used for normalization.

### Identifying differentially expressed micro-RNA or linc-RNA

Simple linear regression analysis was used to associate the expression levels of miRNA and lincRNA to several variables that measured the neuro-pathology in the AD brain. These included either the numbers of neuritic plaques, neurofibrillary tangles or a binary variable representing the pathologic diagnosis of AD on autopsy according to the NIA Reagan criteria [22], All the associations were adjusted for age, sex, study (ROS or MAP), the proportion of neurons in the tissue [2], RNA Integrity number (RIN) and post-mortem interval (PMI). The proportion of neurons in the tissue is estimated from DNA methylation data available from the same brain region of each individual, as described in our recent study [2], A Bonferroni corrected significance threshold of 0.05 was used to account for multiple comparisons.

### Constructing micro-RNA and linc-RNA networks

Linear models were used to identify miRNA or lincRNA that were associated with either NP, NFT or AD. To do this the models included age, sex, study, NNLS, RIN, PMI, NP, NFT and AD as covariates. Using these models a miRNA or lincRNA were included in the networks if there was evidence that any of effects sizes of NP, NFT or AD were non-zero (nominal p-value from an F-test from less than 0.05). For each of the included miRNA or lincRNA forward stepwise variable selection was used with a BIC selection criteria to select which edges between miRNA or lincRNA and explanatory variables should be included in the network. As RIN is associated with all the miRNA and lincRNA its edges are excluded from the networks.

## Declarations

### Ethics approval and consent to participate

The ROS and MAP studies were approved by the Institutional Review Board of Rush University Medical Center. All subjects have given written informed consent.

### Consent for publication

Not applicable

### Availability of data and material

The datasets generated and analyzed during the current study are available on the AMP-AD Knowledge Portal https://www.svnapse.org/#!Svnapse:svn3219045

## Acknowledgements

We would like to thank the participants of the ROS and MAP studies for their participation in these studies. Support for this research was provided by grants from the US National Institutes of Health (R01 AG036042, R01 AG036836, R01 AG17917, RF1 AG15819, R01 AG032990, R01 AG18023, RC2 AG036547, P30 AG10161, U01 AG46152, P50 AG016574, U01 ES017155, K25 AG041906-01) and the Rainwater Foundation/Tau Consortium.

## Author contributions

C.M., A.T., A.M.K. and S.I collected, prepared and generated data from the samples. E.P., J.X., N.P. and S.R. performed analyses on the resulting data. P.L.D. and D.A.B. designed the study. E.P, P.L.D and S.R. wrote the manuscript. All of the authors critically reviewed the manuscript

